# Prediction of Oligomeric Status of Quaternary Protein Structure by using Sequential Minimal Optimization for Support Vector Machine

**DOI:** 10.1101/2024.08.15.608027

**Authors:** Suresh Kumar

## Abstract

The prediction of quaternary structure and its function is important in pharmaceutical applications, drug design, medical or engineering applications. With the drastic rise in the number of protein sequences presented to the public database, acquiring an understanding of possible quaternary structural characteristics of their interested proteins in a timely way is essential for structural genomics research. There is a need for a method to identify quaternary attributes based on its amino acid sequence. In this study, discrimination of hetero-oligomer from non hetero-oligomer was demonstrated using amino acid composition implemented using a support vector machine with a minimum sequential optimization technique.

## Introduction

The technology of high-throughput sequencing (HTS) has dramatically altered the genetic discovery landscape [1]. These technologies have improved the production of a sequencing method by several orders of magnitude, allowing the entire genome to be sequenced and a dozen of exomes per week [2]. Due to the development of sophisticated DNA sequencing technology, the protein sequences were rapidly accumulated in protein sequence databases [3]. It is vital for both biologists and computer scientists to extract these protein sequences information for helpful understanding. With the fast increase in the protein sequence owing to different genomics initiatives, it is important to develop a technique for identifying protein quaternary status from the amino acid sequence using a machine-learning method. The amino acids are the building blocks of proteins. Proteins contain the information necessary to determine how the protein folds into a three-dimensional structure and the stability of the resulting structure in their amino acid sequences [4]. Proteins exhibit a hierarchy of primary, secondary, tertiary and quaternary structural features that guides our current understanding of their biological features and evolutionary origins [5]. The linear sequence of amino acids is regarded as the protein’s primary structure. The secondary structure relates to local folded structures forming within a polypeptide due to backbone atom interactions. Depending on hydrogen bonding, stretches or strands of proteins or peptides have distinct local structural conformations or secondary structure. The alpha helix and beta-sheet are the two primary kinds of secondary structure. An entire protein molecule’s general three-dimensional shape is the tertiary structure. Many proteins are made up of more than one polypetide chains, they are known as quaternary structure. Each polypeptide chain is called as a subunit in protein, this may consist of the same or separate chain. Many proteins tend to self-associate into assemblies of two or more polypetide chains. Protein assemblies composed of one polypeptide chain are called monomers, and more than one polypeptide chain is called oligomers [6]. If the protein complex is made up of identical subunits, it is called a homo-oligomer; otherwise it is called a hetero-oligomer. Classification can be grouped into dimers, trimers, tetramers and so on based on the number of subunits. Although substantial progress has been made in analyzing protein structure with different experimental methods, experimentation is typically costly and time-consuming to determine protein structure [7]. Protein quaternary structure prediction is a significant problem in current bioinformatics. Consequently, it is essential to create a prediction method for protein quaternary assembly states that will allow the study of protein structure and function using the present and rapidly expanding the volume of sequence information. It is usually recognized that most of the proteins contain the necessary information to form in to a function three-dimensional shape. Many proteins consist of more than one polypeptide chain often referred to as protein subunits. The quaternary structure relates to how these protein subunits interact and organize to form a bigger complex of proteins. This indicates that primary sequences contain information about the quaternary structure. Many computational methods are proposed to identify the quaternary protein structure from the amino acid sequence. Garian suggested a technique for predicting homo- and non-homo-dimers using decision tree method [8]. Chou & Cai has created a strong technique by which a specific protein chain can be identified as monomer or homo-oligomer [9]. Zhang et al. created a technique for identifying homo-dimer using machine learning approach [10]. Song and Tang introduced the function of the degree of disagreement to discriminate homo-dimers [11]. Xiao and Lin used the grey dynamic model to determine for predicting quaternary attributes [12]. A method that can discriminate between hetero-oligomer and non hetero-oligomer was proposed in this research using sequential minimum optimization method.

## Materials and Methodology

### Datasets

Based on their quaternary status, amino acid sequences were downloaded from the Uniprot database [13] release 15.6 using the keywords: “monomer,” “homodimer,” “homotrimer,” “homotetramer,” “homopentamer,” ‘homohexamer,” “homoheptamer,” “homo-octamer,” “heterodimer,” “heterotrimer,” “heterotetramer,” “heteropentamers,” “heterohexamer,” “heteroheptamer,” or “heterooctamer. Moreover, the number of heptamer and octamer is too small to be analyzed so it is removed from the dataset construction. The sequences downloaded include 11096 monomers, 43088 homo-oligomers and 13669 hetero-oligomers. Sequences containing amino acids like ‘X’ or ‘Z’ and membrane proteins recognized using the TMHMM server [14] have been omitted. Redundancy of the sequence or homology bias to any other in the same subset was removed by using the CD-HIT algorithm [15] which has greater than 40 percent sequence identity. After filteration, the total number of proteins was reduced to 1404 monomeric, 2982 homo-oligomeric and 1444 hetero-oligomeric proteins with a total of 5830 protein chains.

### Amino acid composition and parameterization of the protein dimension

The composition of amino acids was used to depict the features of the sequence of amino acids. The composition of amino acids is the percentage of all twenty amino acids that occur naturally. It is known that the amino acid composition depends on the protein dimension [16,17]. For this reason, the data sets were categorized into 11 subsets with increasing of 100 residues: 0-100, 100-200,200-300, 300-400, 400-500, 500-600, 600-700, 700-800, 800-900, 900-1000 and above 1000.

### Function-SMO classifier

Many different prediction algorithms have been developed to address this problem. Here, we focused on the Function-SMO which was implemented by using the Waikato Environment for Knowledge Analysis (WEKA) machine learning software [18]. The classifier SMO in Weka is a kind of support vector machine, implementing John Platt’s sequential minimal optimization algorithm. It is an algorithm to fix the quadratic programming problem that happens during support vector machine training. For a comprehensive description of the classifier, readers are referred to as reference [19, 20].

### Statistical assessment of the predictive model

The performance is measured by using a jack knife test. The training dataset is randomly dived into *n*-subsets equally, where *n-1* subsets are used to train the model and the remaining subset is to evaluate the model and it is repeated n times. If is the number of the samples, it was named jack knife test (or leave-one-out cross validation) [21]. 10-fold cross-validation was used here in this research to assess the classifier’s precision. To assess the predictive performance of the suggested model, five measurements were used: sensitivity (Sn), specificity (Sp), accuracy (Acc) and Matthews’ correlation coefficient (MCC) [22]. These parameters are defined as follows:

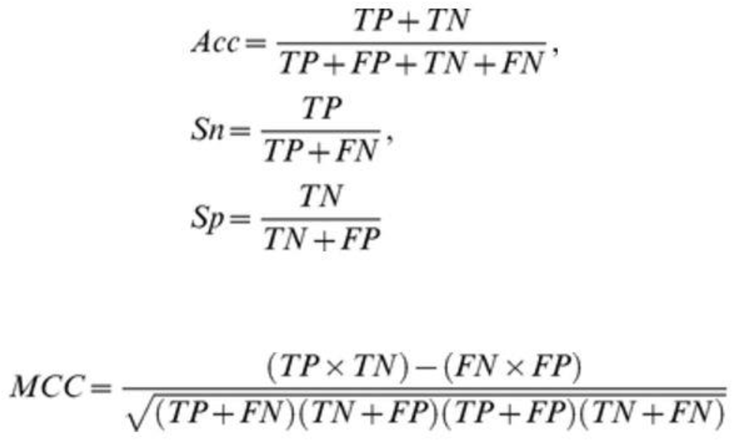

where TP is a true positive number, TN is true negative, FP is false positive, FN is a false negative.

## Results & discussion

In this study, the available protein sequence data available in Uniprot (Table 1) was used to design a computational method to discriminate hetero-oligomer and non-hetero-oligomer. The amino acid composition was used in this study, as it generally accepted which contains all information to fold proper quaternary structure [23]. The amino acid composition was used in various computational methods to predict various levels of protein structure [24-26]. In this study function-SMO classifier was used to discriminate hetero-oligomer from non-heter-oligomer (monomer and homo-oligomer). A Support Vector Machine (SVM) is a classifier which separates the two classes by finding the hyper-plane. SMO uses the polynomial kernel to implement the sequential minimal optimization algorithm to train a support vector classifier. Based on the protein dimension, 11 subgroups from a range of 0-100 to above 1000 amino acid residues were used training the classifier (table 1).

**Table 1:** Number of protein chains within each of the 11 subgroups of protein dimensions.

| *Range of number of residues* | *Monomeric protein chains* | *Homo-oligomeric protein chains* | *Hetero-oligomeric protein chains* |
| --- | --- | --- | --- |
| 0-100 | 111 | 127 | 95 |
| 100-200 | 189 | 381 | 222 |
| 200-300 | 232 | 576 | 208 |
| 300-400 | 238 | 568 | 182 |
| 400-500 | 196 | 457 | 176 |
| 500-600 | 142 | 265 | 113 |
| 600-700 | 92 | 158 | 103 |
| 700-800 | 65 | 98 | 82 |
| 800-900 | 40 | 63 | 66 |
| 900-1000 | 35 | 55 | 45 |
| >1000 | 64 | 234 | 152 |
| Total | 1404 | 2982 | 1444 |

Often three cross-validation techniques are used to determine predictor efficiency are an independent dataset, sub-sampling and jack-knife testing. In the jack-knife cross-validation method, each protein in the dataset identified as a tested protein and remaining proteins will be used for training the predictor. In this study, a 10-fold cross-validation jack knife test was used. In perspective of this, the 10-fold cross-validation jack-knife test was introduced to examine the present method’s predictive quality.

The performance of the algorithms was evaluated using the Matthews Correlation Coefficient (MCC), sensitivity, specificity, and accuracy. The classifier accomplished an overall accuracy of 79.1 percent by discriminating hetero-oligomers from non hetero-oligomers (table 2). From table 2, it is observed that the support vector machine with sequential minimum optimisation i.e function-SMO algorithm resulted in high accuracy (79.1%) while Mathews’s correlation is 0.4 in discriminating hetero-oligomer from non-hetero-oligomer (homo-oligomer & monomer). Also, the prediction accuracy of the classifier based on the protein dimension was measured (figure 1). The accuracy of hetero-oligomeric and non hetero-oligomeric protein discrimination ranged from 74 percent (100-200 residues) to 80 percent (500-600 residues). The highest predictive accuracy (above 80%) is between 300-400 and 500-600 residue proteins. The reason for variation in the classifier’s accuracy is not surprising given Uniprot’s relative dearth that has proper quaternary status annotation [27, 28], and this will decline in the future as the database dimension increases the quaternary status and improves its annotations.

**Table 2:** Prediction performance of the classifiers in discriminating heterooligomeric from non-hetero-oligomer (monomer & homo-oligomer) with 10-fold cross-validation using SVM-SMO method.

| *Classifier* | *Sensitivity (%)* | *Specificity (%)* | *Accuracy (%)* | *Matthews C.C* |
| --- | --- | --- | --- | --- |
| Function-SMO | 70.4 | 76.8 | 79.1 | 0.480 |

**Figure 1:**
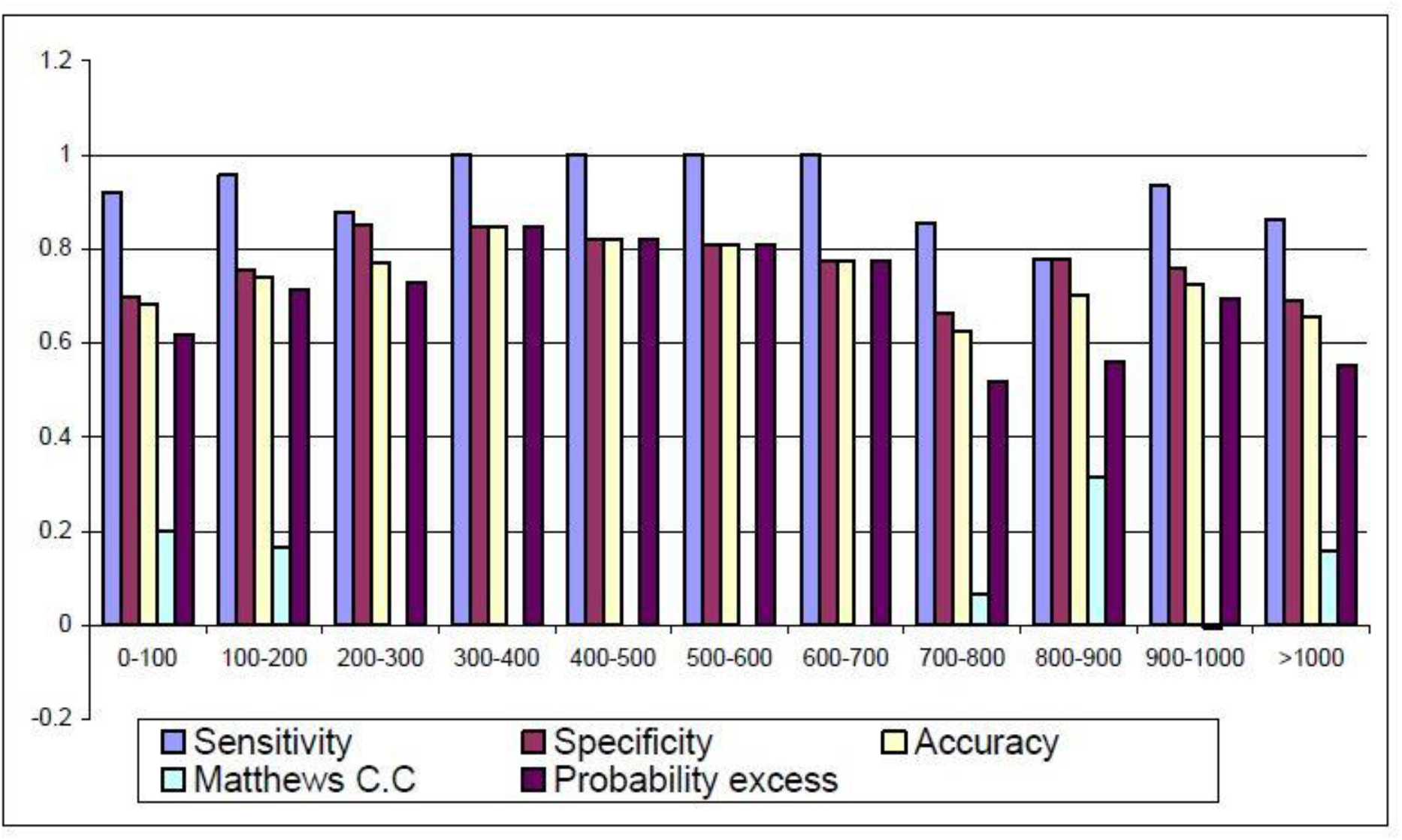
Prediction reliability among 11 subgroups of protein chains of different dimension (function SMO classifier).

## Conclusion

In this study, a support vector machine with a sequential optimization method is adopted to show discrimination of hetero-oligomer and non hetero-oligomer (monomer and homo-oligomer) with high accuracy using amino acid composition. Also, based on the protein dimension study, it is proved the accuracy of hetero-oligomer discrimination from non-hetero-oligomer increases when there is more data available. As more and more information is being added to protein databases in the future, a precise prediction method could be created that would significantly support quaternary status prediction.

## Acknowledgment

The author thanks Management & Science University, Shah Alam, Selangor Darul Ehsan, Malaysia for technical help to carry out this researh.

